# Characterization of Sorbitol Dehydrogenase SmoS from *Sinorhizobium meliloti* 1021

**DOI:** 10.1101/689042

**Authors:** MacLean G. Kohlmeier, Ben A. Bailey-Elkin, Brian L. Mark, Ivan J. Oresnik

## Abstract

*Sinorhizobium meliloti* 1021 is a Gram-negative alphaproteobacterium with a robust capacity for carbohydrate metabolism. The enzymes that facilitate these reactions assist in the survival of the bacterium across a range of environmental niches, and they may also be suitable for use in industrial processes. SmoS is a dehydrogenase that catalyzes the oxidation of the commonly occurring sugar alcohols sorbitol and galactitol into fructose and tagatose respectively using NAD^+^ as a cofactor. The main objective of this study is to evaluate SmoS using biochemical techniques. The nucleotide sequence was codon optimized for heterologous expression in *E. coli* BL21 (DE3) GOLD cells, the protein was subsequently overexpressed and purified. Size exclusion chromatography and X-ray diffraction experiments suggest that SmoS is a tetrameric peptide. SmoS was crystallized to 2.1 Å in the absence of substrate and 2.0 Å in complex with sorbitol. SmoS was characterized kinetically and shown to have a preference for sorbitol despite a higher affinity for galactitol. Computational ligand docking experiments suggest that galactitol oxidation proceeds slowly because tagatose binds the protein in a more energetically favorable complex than fructose, and is retained in the active site for a longer time frame following oxidation which reduces the rate of the reaction. These results supplement the inventory of biomolecules with the potential for industrial applications and enhance our understanding of metabolism in the model organism *S. meliloti*.

## Introduction

Sugar alcohols, also called polyols, are carbohydrate compounds that can be formed by the reduction of an aldo or keto sugar. The first polyols were identified from honeydew, a substance secreted by aphids as they feed on plant sap [1]. The most commonly encountered sugar alcohols in nature are sorbitol, mannitol, and galactitol (also known as dulcitol or melampyrite) [2]. These linear, six carbon polyols were named for the higher plants from which they originated; sorbitol from *Sorbus aucuparia*, mannitol from *Fraxinus ornis* or manna ash, and galactitol from *Melampyrum nemorosum* [1].

Sugar alcohols and their derivatives have a variety of applications. Sorbitol is commonly included in food products for sweetness, texture, and preservation, and can be present in pharmaceuticals [3, 4]. D-tagatose, a product of galactitol oxidation, is classified as a rare sugar and is being considered as a treatment for diabetes due to its insulin independent metabolism in humans and potential to lower blood glucose levels [5–7]. The concentrations of sugar alcohols in plant tissue are typically too low for chemical extractions to generate sufficient yields, therefor polyols are often synthesized for commercial use via catalytic hydrogenation of more readily available sugars [4]. However, biological enzymes can serve as biocatalysts for the generation of sugar alcohols and related molecules at an industrial scale. Some advantages to biocatalysts include high product selectivity and low environmental or physiological toxicity [8]. As an example, galactitol dehydrogenase has been immobilized on gold electrodes for use in electrochemical reactors with the goal of generating precursor molecules for pharmaceuticals via reactions that regenerate reduced cofactors [9–11].

Enzymes of microbial origin are ideal with respect to industrial applications as they can be produced in an easy, cost effective, and consistent manner [12]. Carbohydrate metabolism in plant associated soil bacteria has been studied in great detail due to the involvement of carbon utilization in symbiotic establishment and efficiency [13, 14]. Transport genes responsible for the uptake of sorbitol, mannitol, and galactitol are induced in the rhizosphere [15]. In bacteria, the initial step of sugar alcohol metabolism is often oxidation into a keto sugar, followed by phosphorylation [16]. The root-nodulating bacterium *Sinorhizobium meliloti* has been shown to produce a D-sorbitol specific dehydrogenase (SDH), which uses NAD^+^ as a cofactor [17]. A mutant lacking fructose kinase activity was unable to grow using sorbitol as a sole carbon source, suggesting that fructose is the product of sorbitol oxidation in *S. meliloti* [18]. A mutation to a gene annotated as a putative sorbitol dehydrogenase *smoS* resulted in a strain with the inability to grow on several sugar alcohols, including sorbitol [19], suggesting that *smoS* encodes the SDH protein. *smoS* was first identified as encoding a SDH in *Rhodobacter sphaeroides*, in which it was described as one gene in a novel polyol metabolic operon, as well as a member of the short-chain dehydrogenase/reductase (SDR) family [20]. SDR proteins are typically about 250 amino acids in length and despite having low sequence identity at 20-30%, members of this family share a similar overall three-dimensional structure [21]. Currently there are over 230,000 members of the SDR family in the UniProt database, and a recently devised nomenclature system based on Hidden Markov Models placed SmoS within the SDR196C subfamily [22, 23]. *Rs*SDH is dependent on NAD^+^ as a cofactor and has activity on sorbitol and galactitol (Fig. 1) [20]. Structural studies on the protein in the absence of bound substrate were some of the first structures of a bacterial SDH in the SDR family [24]. The purpose of this study is to characterize SmoS from *S. meliloti* with respect to its structure, as well as kinetic and physical properties.

**Fig 1.**
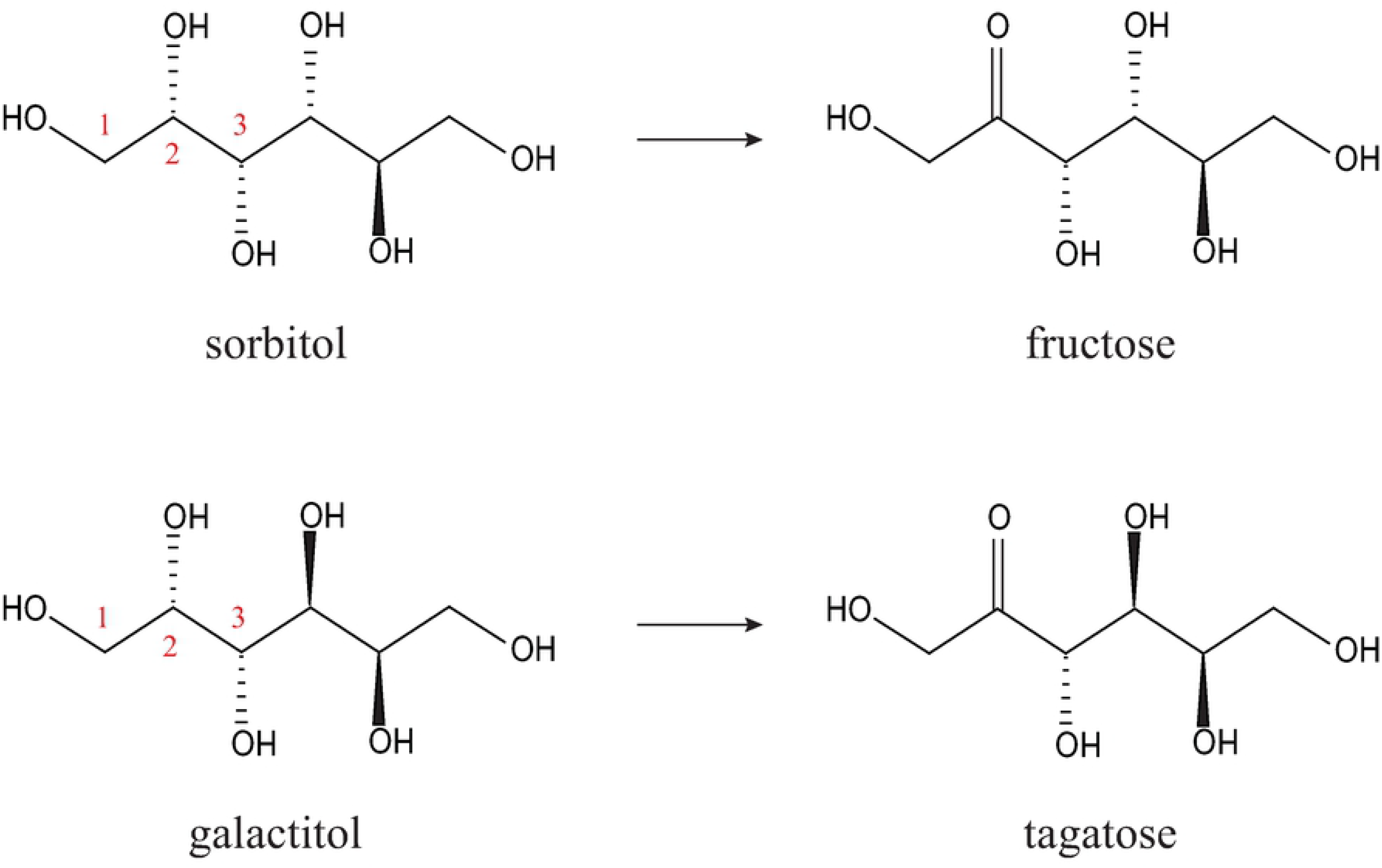
Enzymatic reactions catalyzed by SmoS. Sorbitol or galactitol are oxidized at carbon 2, using NAD^+^ as a cofactor, producing fructose or tagatose respectively, as well as NADH. Both sugar alcohols are viable substrates for SmoS due to the identical orientation of hydroxyl groups about carbons 1, 2, and 3.

## Results

### Structural characterization of *Sm*SmoS

Size exclusion chromatography of purified SmoS showed two distinct peaks at elution volumes of 49.22 mL and 55.35 mL (Fig. 2A), with the most prominent peak at ~55 mL likely representative of a tetrameric complex of SmoS. To corroborate these results, the column fractions were separated by nondenaturing polyacrylamide gel electrophoresis and stained with Coomassie Brilliant Blue. Fractions 2-4 showed two distinct bands, while fractions 5-9 contain a single band, which mimics the migration distance of the lower band from fractions 2-4 (Fig. 2B). Both protein bands are capable of sorbitol oxidation when the gel is stained for dehydrogenase activity (Fig. 2B), and resolve to a molecular weight of 27 kDa when SDS is included in the gel matrix (Fig. 2B), suggesting that both bands observed are due to the presence of SmoS.

**Fig 2.**
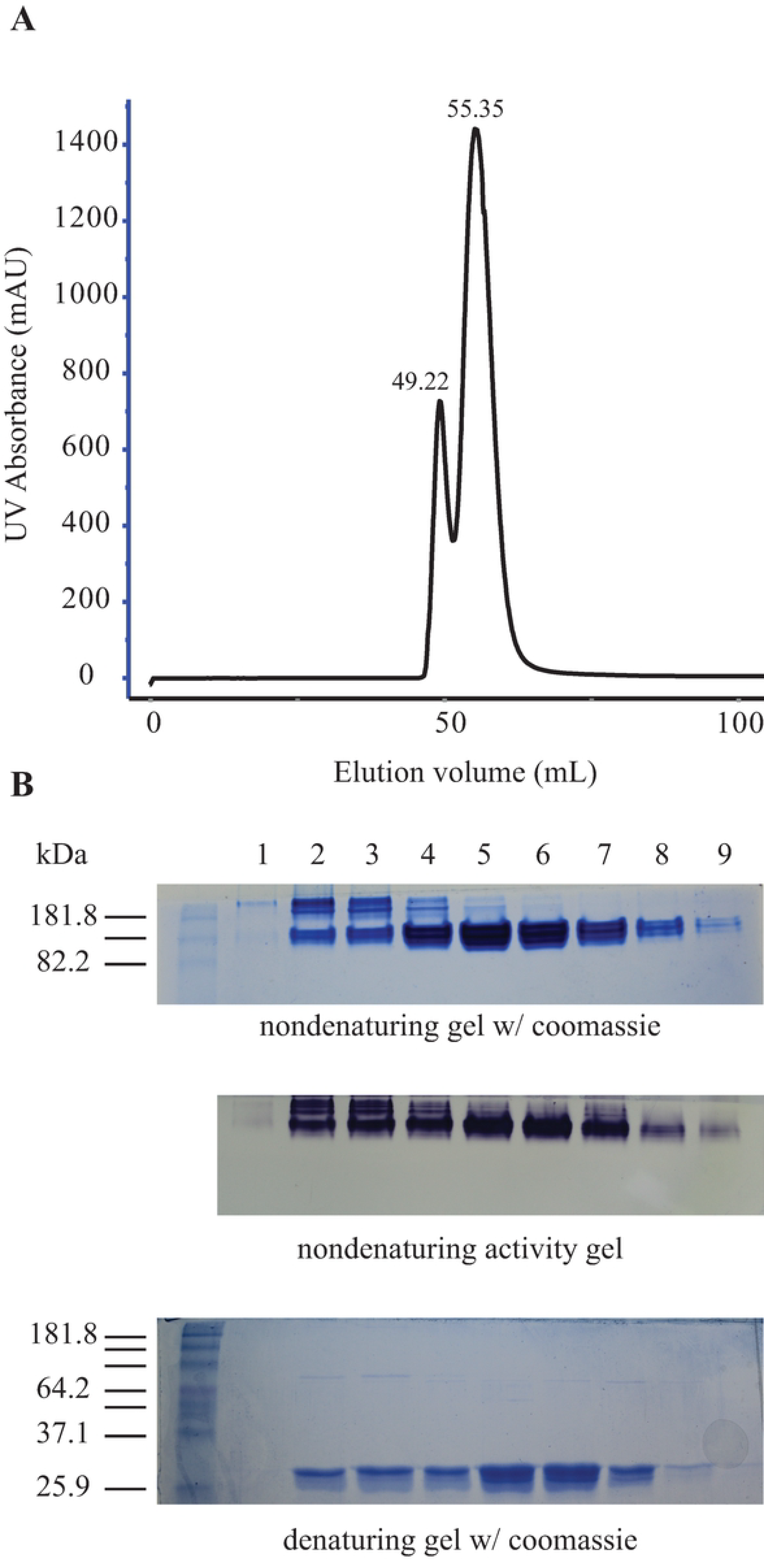
Size exclusion chromatography of purified SmoS from a Superdex 75 gel filtration column and analysis by polyacrylamide gel electrophoresis (PAGE). (A) UV trace of elutions from the S75 column displaying two peaks at approximately 49 mL and 55 mL. (B) Elutions separated by nondenaturing PAGE stained with coomassie blue (top), elutions separated by nondenaturing PAGE stained for sorbitol dehydrogenase activity (middle), and elutions separated denaturing PAGE and stained with coomassie blue (bottom).

To further characterize SmoS, the enzyme was crystallized and determined to a resolution of 2.1Å (Fig. 3; Table 1). Consistent with this observation, SmoS crystallized as a tetramer, with four copies in the asymmetric unit arranged as a dimer of dimers, similarly to a previously determined structure of a *Bradyrhizobium japonicum* D-sorbitol dehydrogenase (Fig. 3A) [25]. These results are consistent with SmoS being present in two distinct conformations in solution, with the majority being tetrameric.

**Table 1.**
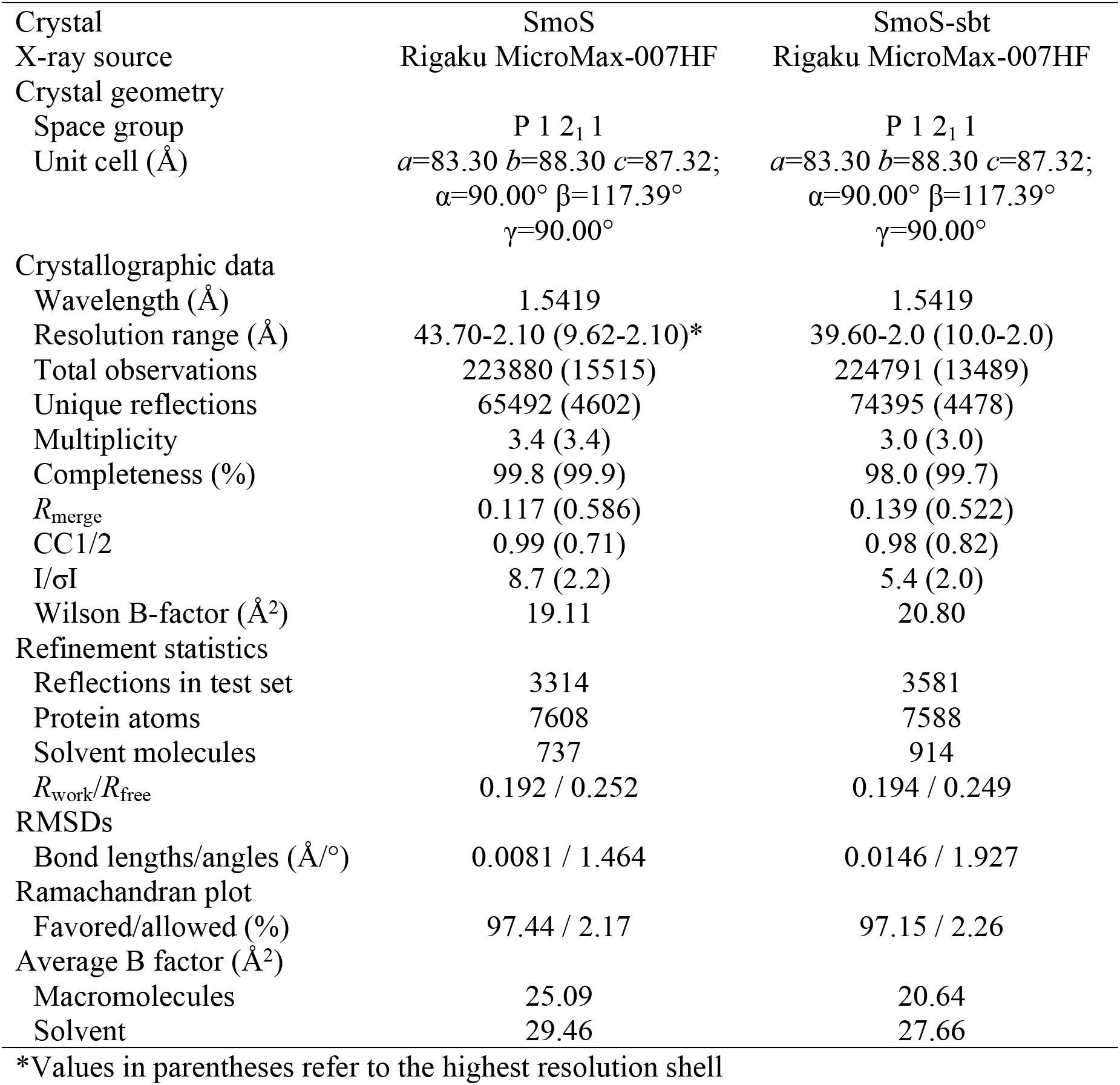
Crystallographic and refinement statistics for SmoS and SmoS-sbt structures

**Fig 3.**
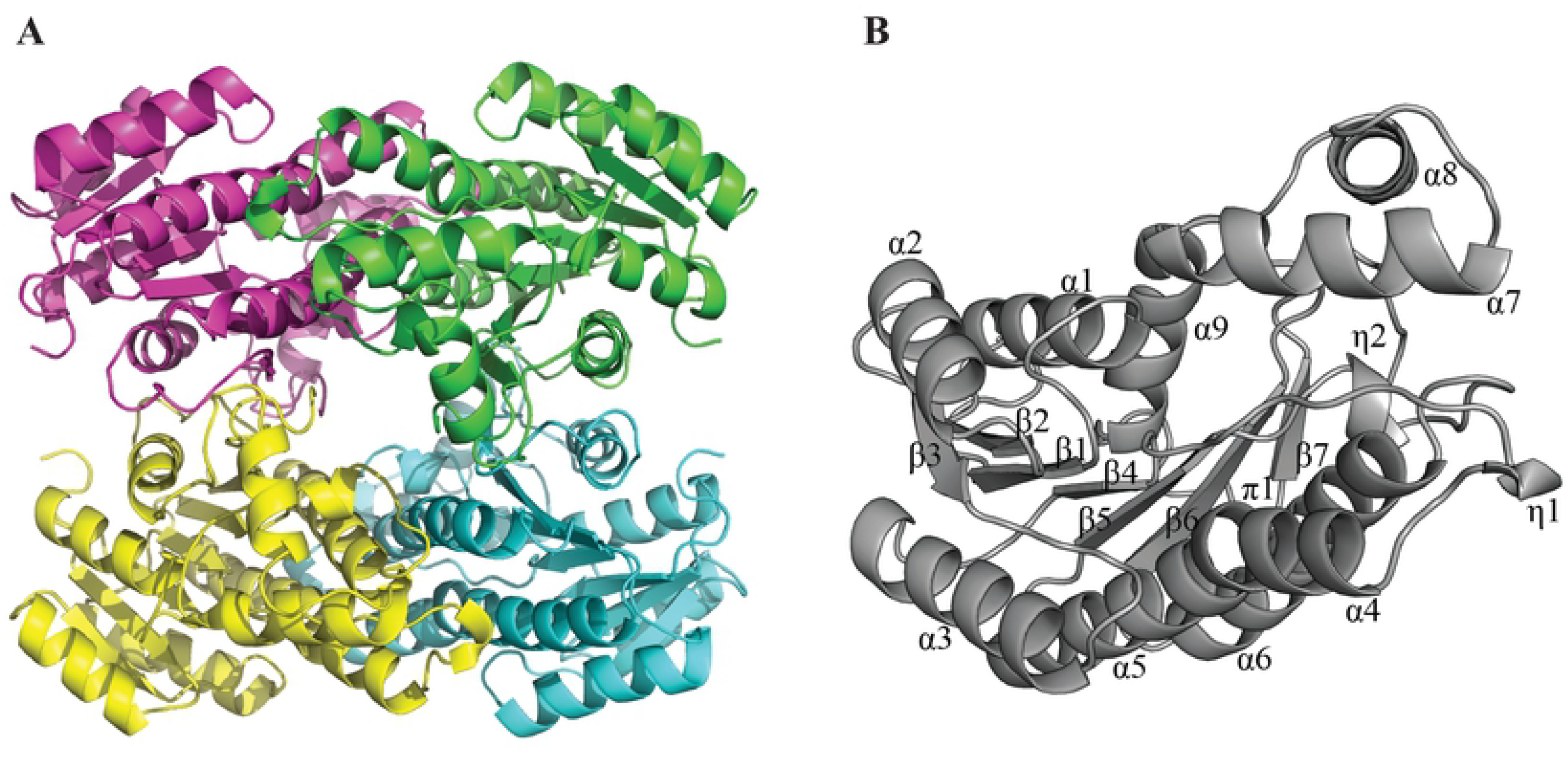
Crystal structure of SmoS from *S. meliloti* 1021. (A) Cartoon representation of the SmoS quaternary structure. SmoS forms a homotetramer; the individual monomers are colored magenta, green, blue, and yellow. (B) Cartoon representation of the SmoS monomer (grey). Secondary structure elements are labeled numerically (α, α-helix; β, β strand; π, π helix; η, 3_10_ helix).

The SmoS monomer adopts a structural fold similar to other previously determined Zn-independent SDR enzymes, comprising of an NAD-binding Rossman fold centralized around a core 7-stranded parallel β-sheet, and an extended α-helical clamp-like lobe formed by helices α7 and α8 involved in substrate binding [24] (Fig. 3B). A DALI search [26] to identify structural homologues of SmoS identified a previously determined *Sm*SmoS structure (deposited by the New York Structural Genomic Consortium), and a *R. sphaeroides* sorbitol dehydrogenase (81% sequence amino acid identity), which aligned to *Sm*SmoS with an RMSD of 0.7 Å over 256 C_α_ atoms, and adopted a nearly identical structural fold [24].

In an attempt to uncover the residues involved in substrate binding, SmoS was also crystallized in the presence of sorbitol and a structure determined to 2.0 Å (Fig.3; Table 1). Consistent with other described Zn-independent SDR enzymes, conserved active site residues Tyr153, Lys157, Ser140 and Asn111 form the active site (Fig. 4A). Residue Asn111 resides on a π-bulge motif formed by an atypical backbone hydrogen bond disrupting helix α4. This deformation allows the backbone carbonyl group of Asn111 to form a hydrogen bond with a water molecule likely to be involved in the formation of a proton relay system similar to what has been described for the *Comamonas testosterone* hydroxysteroid dehydrogenase [27, 28]. Clear electron density representing sorbitol was visible near the active site of each of the four monomers in the asymmetric unit, with sorbitol coordinated near the active site through a hydrogen-bonding network mediated by SmoS residues Gln141, Glu147, Gly184 and His190 (Fig. 4A and B). A comparison of the apo and sorbitol-bound forms of SmoS reveals a slight change in the position of the clamp domain formed by helices α7 and α8, which moves inward during sorbitol binding and allows for the satisfaction of a hydrogen bond between His190 and sorbitol OH1 (Fig. 4C). Interestingly, while clear density for sorbitol was observed in all *Sm*SmoS monomers, the substrate does not appear to be positioned appropriately within in the active site to permit NAD^+^-mediated oxidation at C2. In order for the reaction mechanism to proceed as described, the sorbitol C2 hydroxyl group would need to be positioned within hydrogen bonding distance from Tyr153, to allow for Tyr153-mediated proton abstraction and subsequent oxidation of C2 *via* the nicotinamide moiety of NAD^+^. In the SmoS-sbt structure, the C2 hydroxyl group is situated ~5.9 Å away from Tyr153, and points away from the active site residue in an arrangement that would not permit the conversion of sorbitol to fructose.

**Fig 4.**
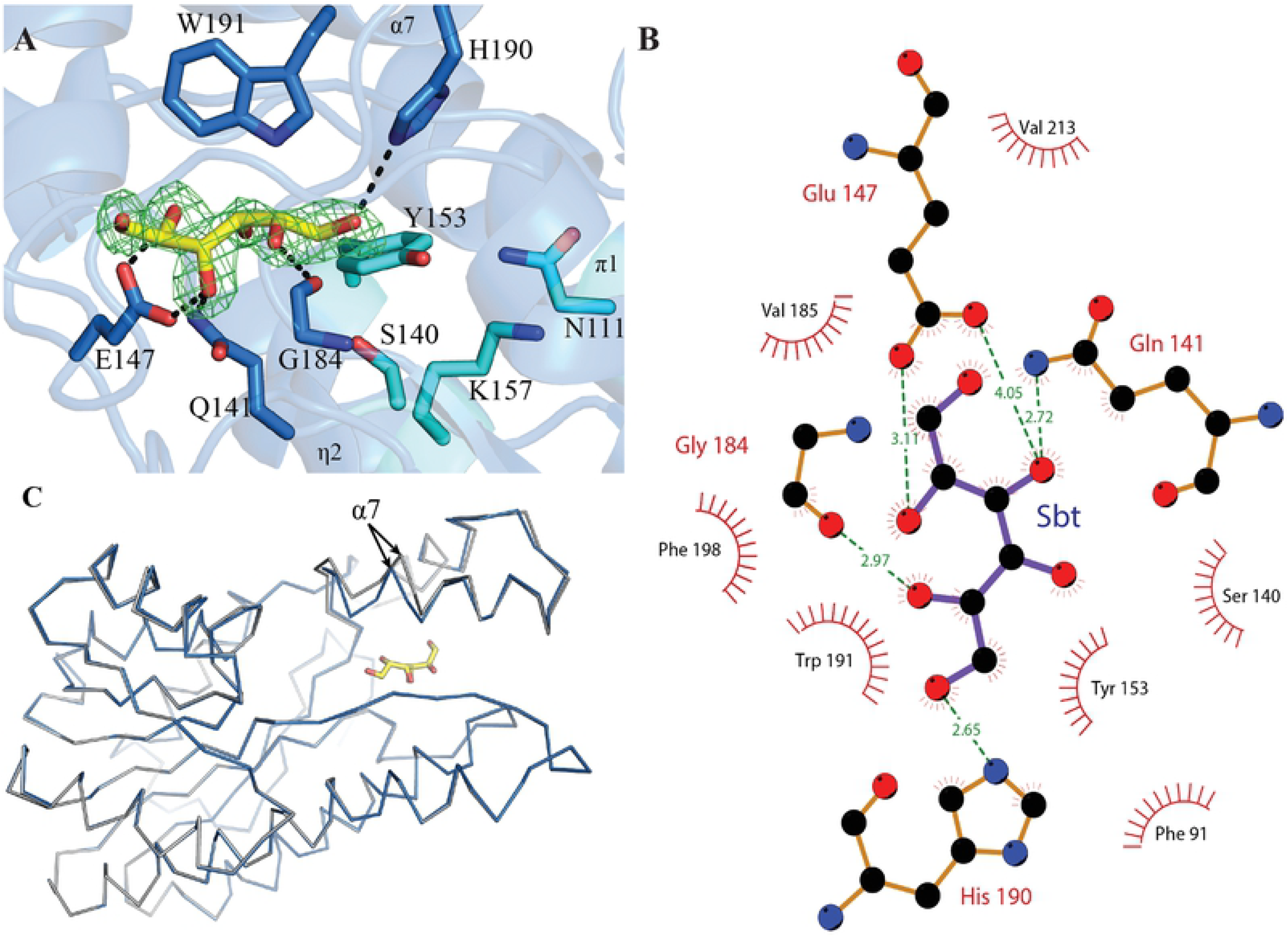
Crystal structure of the SmoS-sbt complex. (A) Close up on the active site of sorbitol-bound SmoS. Catalytic residues are shown as blue sticks, and residues involved in the coordination of sorbitol are shown as a cyan sticks. Sorbitol is shown as yellow sticks, surrounded by an *mF*_o_-*DF*_c_ omit map generated using phenix.polder ([29]; green mesh) contoured to 3.0σ. (B) Two-dimensional representation of the H-bonding network observed in the SmoS-sbt complex. Carbon atoms are black, oxygen atoms are red, nitrogen atoms are blue, H-bonds are shown as green dashed lines with corresponding bond lengths (Å). Figure was generated using LigPlot [30]. (C) Superposition of apo SmoS (grey), and SmoS-sbt (blue) depicted in ribbon diagrams with the movement of helix α7 indicated by arrows.

### SmoS has a high pH optimum and a preference for sorbitol

It has been reported that functionally related enzymes to SmoS have optimum activity at alkaline pH levels [31, 32]. To investigate the pH preference of *S. meliloti* SmoS, sorbitol dehydrogenase assays were conducted across a pH gradient facilitated by several solutions of differed buffering capacities. 1 µg of SmoS was added to the assay mixture along with 10 mM sorbitol and 1.5 mM NAD^+^, the buffers included MES, MOPS, TRIS, and CAPS, each at a concentration of 20 mM, which allowed for a pH gradient spanning pH 5.5-12.5. An optimum enzyme activity of 57.8 mM/min/mg was observed at pH 11; activities recorded across the gradient are reported relative to this value (Fig. 5). Fifty percent of this activity was found at pH 9.5. All subsequent activity assays were conducted in a solution buffered with 20 mM CAPS pH 11. This result is consistent with observations made in *R. sphaeroides* [33].

**Fig 5.**
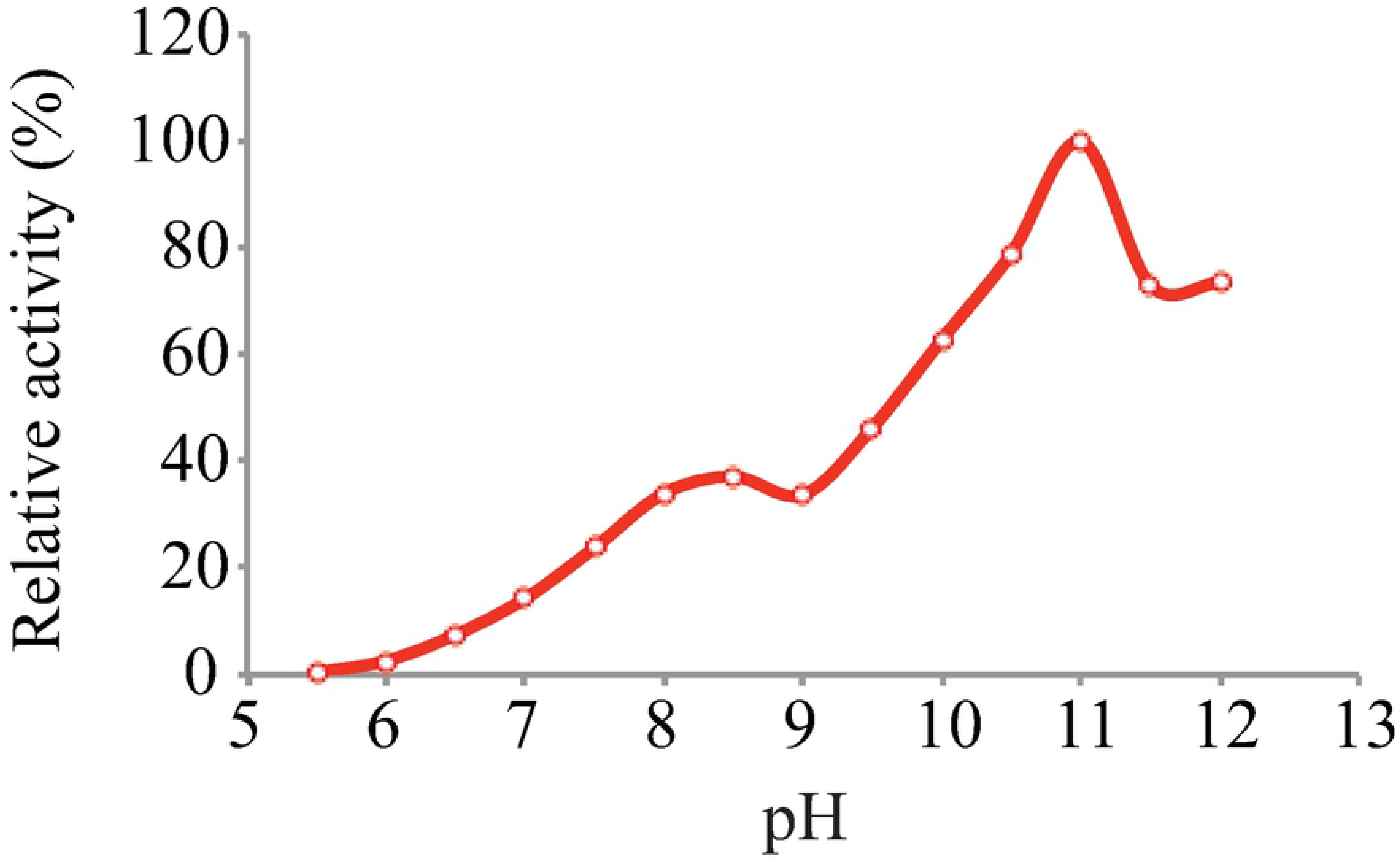
Effect of pH on SmoS dehydrogenase activity. Reactions were carried out with10 mM sorbitol using 200 mM MES, MOPS, TRIS, or CAPS buffers over their appropriate pH ranges. Activity at the optimal pH was defined as 100%.

Despite the previous inability to detect galactitol dehydrogenase activity [17], recent work has shown that *S. meliloti* is capable of galactitol oxidation and that SmoS is responsible for this activity [34]. The ability of SmoS to oxidize sorbitol and galactitol is likely due to the stereochemistry of the functional groups about carbon’s 1, 2, and 3, which are identical for both substrates (Fig. 1). To determine the substrate preference of the enzyme, reaction rates were determined by measurement of NADH accumulation over time in a spectrophotometer at 340 nm. Saturation curves for sorbitol and galactitol dehydrogenase activities were generated along with double reciprocal plots facilitating the determination of Michaelis-Menten reaction constants (Fig. 6). It was determined that SmoS has a K_M_ of 2.5 mM for sorbitol, and a K_M_ of 1.2 mM for galactitol (Table 2), however, the maximum velocity (V_max_) of the sorbitol oxidation reaction was calculated to be 50.8 mM/min, while galactitol oxidation proceeded at only 6.4 mM/min (Table 2). Despite a higher affinity, the low reaction velocity of galactitol oxidation greatly reduces the overall reaction efficiency (k_cat_/K_M_). We note that L-iditol shares hydroxyl group orientation about carbons 1, 2, and 3, with sorbitol and galactiol, however this substrate was not tested due to lack of availability [24].

**Table 2.** Kinetic properties of *Sm*SmoS

| Substrate | $K_M$ (mM) | $V_{max}$ (mM/min) | $k_{cat}$ ($s^{-1}$) | $V_{max}/K_M$ ($min^{-1}$) | $k_{cat}/K_M$ ($mM^{-1}s^{-1}$) |
| --- | --- | --- | --- | --- | --- |
| Sorbitol | 2.5 | 50.8 | 25625.6 | 20.6 | 10419.7 |
| Galactitol | 1.2 | 6.4 | 3227.1 | 5.2 | 2638.9 |

**Fig 6.**
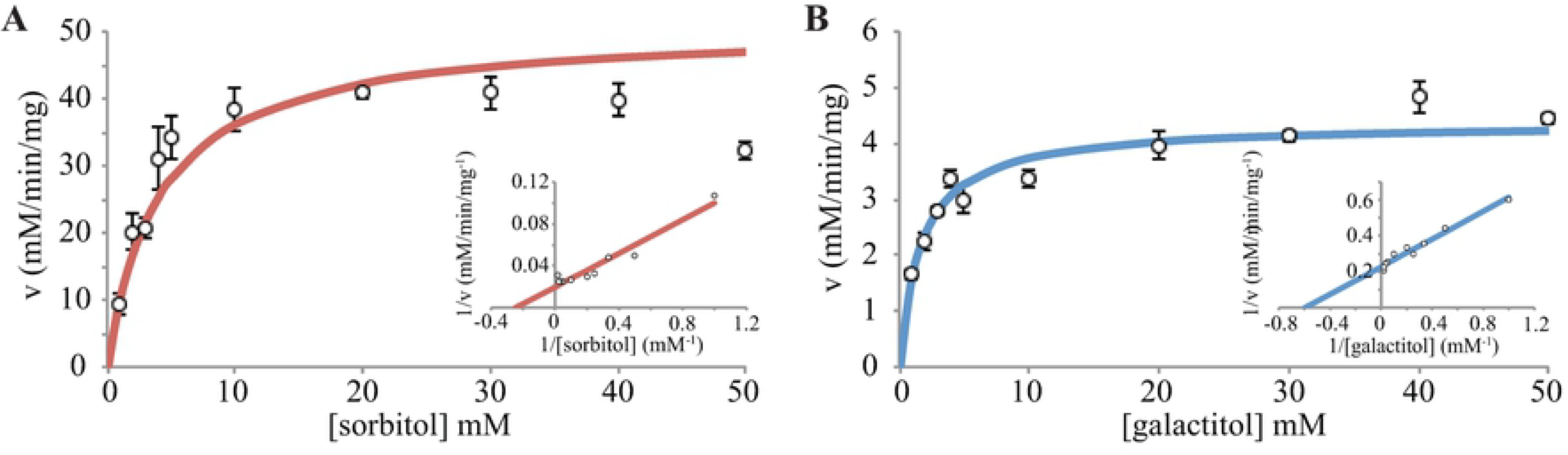
Kinetic characteristics of analysis of *Sm*SmoS. Analysis of sorbitol (A) and galactitol (B) oxidation by using Michaelis-Menten and Lineweaver-Burk plots.

### SmoS-tagatose complex is predicted to be in a lower energy state than SmoS-fructose complex

Kinetic analysis revealed that galactitol turnover is much less efficient than sorbitol oxidation (Fig. 6, Table 2). This observation was particularly interesting due to the K_M_ value of galactitol oxidation, which suggested that the enzyme’s affinity for galactitol was higher than for sorbitol (Table 2). This led to the hypothesis that tagatose is a poor leaving group in comparison to fructose and the inability of tagatose to quickly leave the active site results in low reaction turnover. This hypothesis is supported by our inability to detect fructose in the active site of SmoS structures determined from crystals grown in the presence of a large concentration (20%) of fructose. To test this hypothesis, computational ligand docking analysis was conducted using the Rosetta Ligand Docking Protocol on the ROSIE server [35–38]. D-fructose and D-tagatose model files were submitted to the ligand dock protocol along with apo SmoS monomer structure, and the outputs were analyzed for indications of the energy state of the complexes. The server generated 200 docking predictions for each SmoS-ligand complex, which were organized via their interface delta scores. The interface delta score represents the total energy of the complex in isolation subtracted from the total energy of the complex with the substrate bound [39]. The ten models with the lowest interface delta score from each complex were selected. The scores from the SmoS-fru model complexes were consistently higher than the scores reported for the SmoS-tag complexes, suggesting that the SmoS-tag complex is in a lower energy state with higher stability than the SmoS-fru complex (Fig. 7A). The data from each SmoS-ligand complex were analyzed for significance via a students *t* test, revealing a *P* value of 1.3×10^−6^. The entire process from submission to the server through data collection and analysis was repeated independently to evaluate reproducibility; the SmoS-tag complexes were consistently in a lower energy state than the SmoS-fructose complexes. The *P* value for the second trial was 2.4×10^−6^. An examination of the hydrogen bonding interactions that mediate binding reveals that the SmoS-tag complex forms an additional hydrogen bond that is not present in the SmoS-fru complex, which further stabilizes the tagatose bound structure (Fig. 7B and C). These data suggest that the SmoS-tag complex is a lower energy and more stable complex than the SmoS-fru complex, and that the predicted interface energies from the SmoS-fru complexes and the SmoS-tag complexes are statistically different. They also support the hypothesis that tagatose is a poor leaving group in comparison with fructose and are consistent with observations of the kinetic properties of the enzyme.

**Fig 7.**
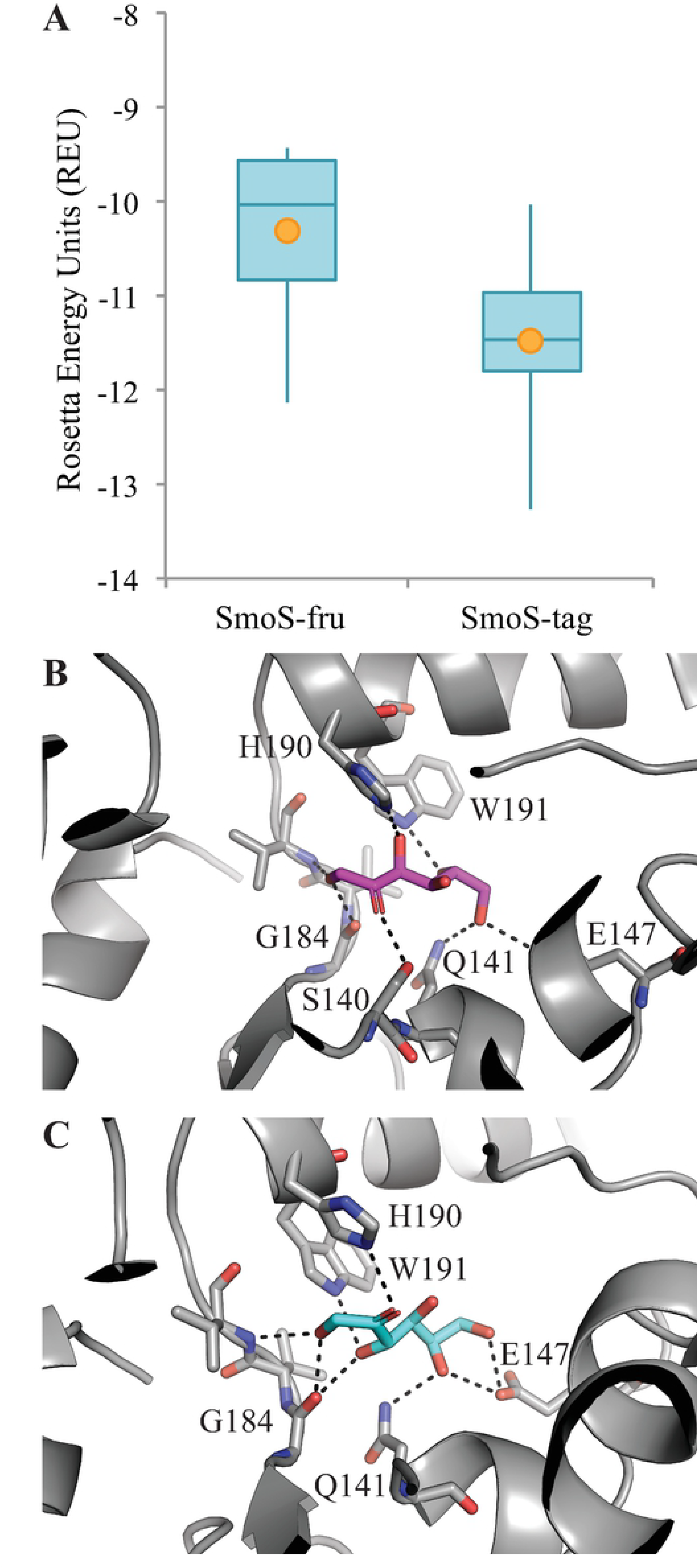
SmoS-fructose and SmoS-tagatose binding complexes predicted by the Ligand docking protocol housed on the ROSIE server. (A) The distribution of the top ten interface delta scores displayed as box and whisker plots. The tips of the whiskers represent the maximum and minimum values, the horizontal lines represent the first, second, and third quartiles, and the orange dots represent the averages of the data sets. P value of 1.3×10^−6^. The lowest energy docking prediction for the SmoS-fructose complex (B) and the SmoS-tagatose complex (C). Fructose is shown in magenta and tagatose in blue.

## Discussion

*S. meliloti* SmoS appears to be most similar to the sorbitol dehydrogenase from *R. sphaeroides*, these enzymes share kinetic characteristics [33], operon structures [20], pH preferences [33], and overall quaternary structure [24]. SmoS can be classified within a group of “high-alkaline enzymes,” which are enzymes with a pH optimum from pH 10-11. These enzymes are useful in industry due to their high durability [40]. Similar to *Rs*SmoS, *Sm*SmoS was found to have a higher affinity toward galactitol compared to sorbitol, but turned over sorbitol at a faster rate [33].

Most of the crystal structures of SmoS related enzymes have reported tetrameric structures found in the crystal packing [24, 25, 41], however reports differ on the structure of the enzyme in solution. *R. sphaeroides* SmoS has been reported as dimeric in solution, on the basis of gel filtration chromatography as well as sucrose gradient centrifugation experiments [33]. However the enzyme was later predicted to function as a tetramer based on predicted surface area exposure [24], and these results were supported by size exclusion chromatography and light scattering experiments [41]. *Bj*SDH had been proposed to exist as a trimer in solution [42] but researchers later suggested that a tetramer was more likely [25]. A galactitol dehydrogenase from *Rhizobium leguminosarum* 3841 has also been reported to be tetrameric in solution [31]. The data presented clearly shows that SmoS from *S. meliloti* is present as a tetramer in solution but with a small subset seemingly present as a hexamer or an octamer made up of a dimer of tetramers (Fig. 2). Of note, it appears that both the tetrameric as well as the higher oligomeric forms show sorbitol dehydrogenase activity (Fig. 2). Tetrameric configurations are reported most often and likely represent the majority of SDR protein structures in solution [43].

The SmoS-sbt structure shows that the hydroxyl group bonded to C1 of sorbitol associating with catalytic residue Tyr153, and that the structure has a subtle difference from the apo structure in that residues His190 and Trp191 in alpha helix 7 are contorted slightly to accommodate the presence of the substrate (Fig. 4C). As well, residues Asn111, Ser140, Tyr153, and Lys157, which have been proposed to be involved in electron transfer, are too distant from the substrate for catalysis (Fig. 4A).

If the positioning of Tyr153 were correct, it would imply that sorbitol should be oxidized to glucose. Based on the available genetic and physiological data it is clear that both sorbitol and galactitol catabolism mediated by SmoS generate fructose and tagatose via an enzymatic reaction in which the hydroxyl group on C2 of the substrate is oxidized forming a planar carbonyl carbon [18, 34]. We also note that enzymes catalyzing the oxidation of sorbitol into glucose are known as sorbitol oxidase (SOX) proteins [44, 45]. These enzymes are dissimilar to SDH enzymes of the SDR family [46, 47].

This anomaly could be due to the absence of NAD^+^ in the binding pocket. NAD^+^ was left out of the crystallization solution because its presence would result in an enzymatic reaction, which would prevent the capture of a substrate-bound complex. However, SDR reactions proceed with the coenzyme binding first and leaving last [48], which may help to explain not only why sorbitol is found in an atypical position, but also why fructose was not found in the active site of the fructose grown crystal structures despite its presence at high concentrations. In addition, modeling of NAD^+^ and sorbitol into the *R. sphaeroides* predicted direct contact and a sandwiching of the C2 carbon of sorbitol between the active site tyrosine, and the nicotinamide ring. Taken together these may explain the observed structure.

Thermal stability of an enzyme can affect its ability to be exploited in industrial processes [8]. It has been proposed that the increased thermal stability of SDH is due to the abundance of proline residues and the proline to glycine ratio in its primary amino acid sequence [25]. Proline is a rigid residue with low configurational entropy due to its pyrrolidine ring hindrance, there are several studies that suggest protein thermostability can be influenced by proline content [49–51]. *Rs*SDH contains 6 proline residues and a Pro/Gly ratio of 0.22, while *Bj*SDH has 13 prolines with a ratio of 0.86. The melting temperatures were found to be 62°C and 47°C respectively (25, 39). The SmoS from *S. meliloti* has 5 proline residues and the Pro/Gly ratio is 0.2, additionally the position of the residues appears to be conserved, indicating that it’s thermostability is likely more similar to *Rs*SDH (Fig. 8).

**Fig 8.**
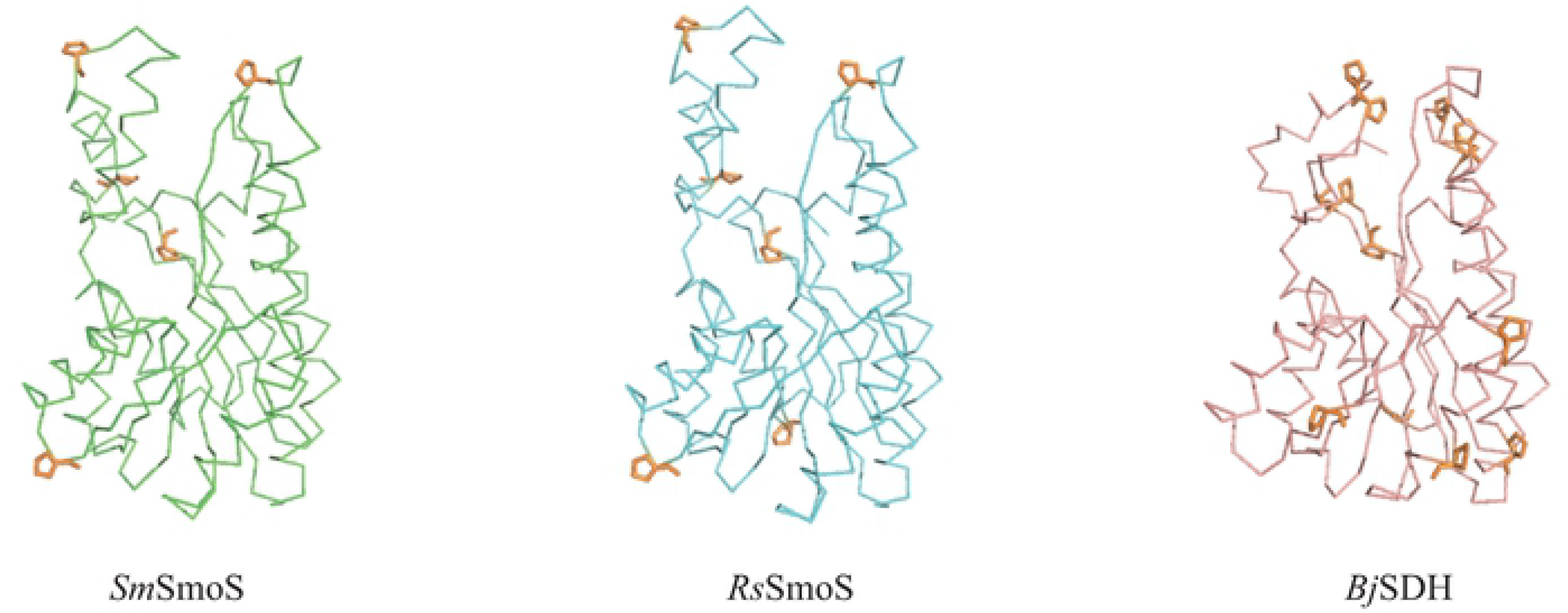
Comparison of the position and distribution of proline residues. *S. meliloti* SmoS (green), *R. sphaeroides* SmoS (blue; PDB ID: 1K2W), and *B. japonicum* SDH (pink; PDB ID: 5JO9), proline residues are shown in orange.

The structure and characterization of *S. meliloti* SmoS provides a high quality structure with sorbitol within the active site. In addition, the characterization and determination of its affinities for its substrates provides insight into why the growth rate of the organism on what should be two equivalent carbon substrates shows great differences. This information is invaluable for higher order resolution of metabolism in *S. meliloti*.

## Experimental procedures

### Bacterial strains and culture conditions

*E. coli* BL21 (DE3) GOLD cells were grown on Luria Bertani (LB) medium [52] at 37°C; when necessary, kanamycin was added to a final concentration of 10 µg/mL in liquid media.

### Overexpression and purification of SmoS

*S. meliloti smoS* is a 774 bp gene with a GC content of 64.5%, the overall GC content of *E. coli* K-12 is 50.8% [53]. To accommodate this disparity, the *smoS* nucleotide sequence was codon optimized for expression in *E. coli*. Translation of *smoS* is predicted to generate a 257 amino acid sequence with a molecular weight of 27.2 kDa [54]. *smoS* was cloned into overexpression vector pET-28a as a *Bam*HI-*Hind*III fragment (GenScript, Piscataway, NJ, USA) and this construct was transformed into competent *E. coli* BL21 (DE3) GOLD cells.

Cultures were grown in 1 L volumes of LB medium at 37°C to an OD_600_ of ~0.6. Induction with 1 mM isopropyl-β-D-galactopyranoside (IPTG) preceded growth overnight, shaking, at 16°C. Cells were pelleted by centrifugation at 10000 rpm for 10 min and stored at −80°C. Pellets were resuspended in 30 mL cold lysis buffer consisting of 50 mM Tris pH 8.0, 300 mM NaCl, 2 mM dithiothreitol (DTT), 10 mM imidazole, and lysed by French Press. Cell debris were removed from extracts by centrifugation at 12000 rpm for 1 hour at 4°C. The cell free lysate was applied to a nickel nitrilotriacetic acid (Ni-NTA) column, which was washed with 10 column volumes of lysis buffer and followed by a second wash with 10 column volumes of lysis buffer with 25 mM imidazole. Final elution was prompted by washing with 3 column volumes of buffer with 500 mM imidazole. Eluted protein was dialyzed against 20 mM HEPES pH 7.5, 150 mM NaCl, 10% (v/v) glycerol, and further purified by gel filtration through a Superdex 75 gel filtration column.

### SmoS crystallization

Purified SmoS was concentrated to 10 mg/mL and screened by sitting drop vapour diffusion using a Gryphon (Art Robbins Instruments, Sunnyvale, CA, USA) robotic drop setter. Screening was performed using 600 nL drops containing SmoS and crystallization solution at a 1:1 ratio, equilibrated against 50 μL of reservoir solution. Initial crystallization hits were identified in 100 mM HEPES pH 7.4, 50 mM sodium acetate and 20% PEG 3000, and further optimized by hanging-drop vapour diffusion using 48-well VDX plates. Crystals of apo-SmoS were grown in 100 mM HEPES pH 7.4, 50 mM sodium acetate and 18% PEG 3000 and crystals of the SmoS-sorbitol complex were grown under the same conditions supplemented with 20% sorbitol, galactitol, tagatose, or fructose. Crystallization with galactitol and tagatose was not pursued due to poor solubility or lack of availability of these respective substrates. Crystals in which sorbitol or fructose were included in the reservoir solution were morphologically indistinguishable from the native crystals.

### X-ray data collection and structure solution

X-ray data for individual SmoS crystals were collected at 100K on a Rigaku MicroMax 007-HF equipped with a RAXIS IV++ detector. X-ray diffraction images were integrated and scaled using XDS [55], and merged using Aimless [56]. Initial phase estimates for apo-SmoS were determined by molecular replacement within Phaser using the deposited structure of *S. meliloti* SmoS (PDB ID: 4E6P) as a search model, and phase estimates for the SmoS-sbt complex were determined using the refined apo-SmoS structure. Structure refinement and model building were performed using REFMAC [57] and Coot [58], respectively within the ccp4i2 software package. All structure figures were generated using PyMOL [59]. The coordinates and structure factors for the apo SmoS and SmoS-sbt structures have been deposited to the Protein Data Bank under PDB ID 6PEI and 6PEJ, respectively.

### Enzyme assays

Spectrophotometric dehydrogenase assays were conducted by measuring the reduction of NAD^+^ at 340 nm for 60 seconds. Reaction buffer consisted of 20 mM CAPS pH 11, 1.5 mM NAD^+^, and increasing concentrations of sorbitol or galactitol, in a total volume of 1 mL. 1µg SmoS was added per reaction. The optimum pH for enzyme activity was determined by measuring dehydrogenase activity using 200 mM MES, MOPS, TRIS, or CAPS to buffer the reaction mixtures over their appropriate pH ranges. All pH-profiling reactions were initiated with 10 mM sorbitol. Additionally, native gel dehydrogenase assays were performed as previously described [60]. Following elution from the S75 column, fractions were separated by nondenaturing polyacrylamide gel electrophoresis; subsequently the gels were stained for dehydrogenase activity with an assay reagent containing Tris pH 8.0, phenazine methosulfate, nitroblue tetrazolium, NAD^+^, and sorbitol.

### Ligand docking analysis

D-fructose and D-tagatose model files, in SDF file format, were submitted to the Ligand Docking Protocol on the ROSIE server, found at http://rosie.rosettacommons.org, along with the apo SmoS monomer structure in PDB file format. The ligand SDF files were downloaded from Research Collaboratory for Structural Bioinformatics Protein Data Bank (RCSB PDB) at https://www.rcsb.org. These ligand models were manipulated to within 5 Å of the SmoS substrate binding pocket using PyMOL [59] prior to submission to add coordinate data to the files. The top ten predicted models with the lowest interface delta scores were collected and the distribution of these data sets was compared with box and whisker plots. A paired *t* test was performed on the score arrays, a *P* value of less that 0.01 was considered significant.

